# Structural Role of Stomatin in Organizing Functional Membrane Microdomains

**DOI:** 10.1101/2025.08.30.673307

**Authors:** Lu Yan, Xinyue Zhou, Meiqi Li, Chenxi Wang, Bailong Xiao, Peng Xi, Peng Zou, Ning Gao

**Author notes:** Correspondence (N.G.); (P. Z.). These authors contributed equally (Lu Yan, Xinyue Zhou).

## Abstract

Stomatin, originally discovered in red blood cells, is a member of the SPFH (Stomatin, Prohibitin, Flotillin, and HflK/C) protein family, which has long been proposed to scaffold functional membrane microdomains (FMMs) enriched in saturated lipids such as cholesterol and sphingomyelin. Stomatin has been reported to associate with a variety of proteins involved in diverse physiological processes, including ion channel regulation, membrane fusion, mechanosensory regulation and vesicle trafficking; however, the mechanisms by which it modulates these interactions remain poorly understood.

Here, we determined the cryo-electron microscopy (cryo-EM) structures of stomatin, revealing its hexadecameric assemblies. The SPFH1 domains insert into the cytosolic leaflet of membranes and, together with the N-terminal hairpin, form a potential cholesterol-binding pocket. Liposome reconstitution experiments demonstrated that stomatin self-organizes into clusters, and Spectrum and Polarization Optical Tomography (SPOT) further showed that these clusters enhance membrane lipid order, supporting the proposed role of stomatin in organizing functional membrane microdomains (FMMs). Proteomic mass spectrometry analysis identified numerous stomatin-associated cargo proteins, including solute carrier (SLC) transporters, RAB GTPases, and integrins, suggesting that stomatin regulates solute transport and contributes to vesicle trafficking and cell migration. Together, these findings elucidate the structural and functional roles of stomatin and underscore its potential as a therapeutic target for modulating cancer cell migration and metastasis.

## Introduction

Erythrocytes of patients with Overhydrated Hereditary Stomatocytosis (OHSt) were first reported to lack a major membrane protein migrating near band 7 on SDS-PAGE, which was initially termed band 7^1–3^ and later renamed stomatin, a name that reflects the markedly increased proportion of stomatocytic erythrocytes in affected individuals^4,5^.

Subsequent studies revealed that stomatin is ubiquitously expressed in diverse tissues and cell types^2,6^, where it participates in multiple physiological processes. Its role in ion channel regulation was first suggested by the elevated permeability of OHSt erythrocytes to monovalent cations^5,7^. Later, stomatin was shown to colocalize with the acid-sensing ion channel ASIC3 (a member of the DEG/ENaC family) and inhibit its activity^8–11^, while also suppressing the ionic current of the gap junction protein pannexin-1 (Panx1)^12^. In addition, stomatin has been implicated in membrane fusion: its depletion in osteoblasts reduces expression of the fusion protein DC-STAMP, impairing osteoclast multinucleation^13^, whereas its overexpression in adipocytes promotes lipid droplet fusion and growth^14^.

In erythrocytes, stomatin also contributes to tethering the membrane skeleton to the plasma membrane (PM), physically associating with glucose transporter 1 (GLUT1), anion exchanger 1 (AE1/SLC4A1), and aquaporin-1 (AQP1)^15^. Moreover, stomatin downregulates GLUT1-mediated glucose transport^16,17^. Beyond red blood cells, stomatin-like protein 3 (STOML3), its closest homolog, is selectively expressed in the nervous system and plays a role in mechanosensation, likely by sensitizing PIEZO1/2 channels or cooperating with stomatin to modulate ASIC3 activity^18–20^.

Despite these documented roles, the molecular mechanisms by which stomatin exerts its regulatory functions remain incompletely understood. As a member of the SPFH (stomatin, prohibitin, flotillin, HflK/C) protein family^21^, stomatin localizes to detergent-resistant membranes (DRMs) enriched in cholesterol and sphingolipids^22–26^, suggesting a role in organizing functional membrane microdomains (FMMs)^24,27^. However, direct structural and mechanistic evidence has been lacking.

In this study, we report the cryo-electron microscopy (cryo-EM) structures of stomatin, revealing its hexadecameric assemblies. Using liposome reconstitution and Spectrum and Polarization Optical Tomography (SPOT), we show that stomatin clusters enhance membrane lipid order, thereby supporting its role in organizing FMMs. Proteomic analysis further identifies solute carrier (SLC) transporters, RAB GTPases, and integrins as key stomatin-associated cargos, linking stomatin to solute transport, vesicle trafficking, and cell migration.

## Results

### 1. Stomatin proteins assemble into cage-like hexadecamers with a variable basal diameter

We employed the human cell line HEK293F to overexpress stomatin (STOM) fused at its C terminus with a Twin-Strep tag, followed by membrane solubilization with DDM and subsequent purification. Negative-staining EM (nsEM) revealed that DDM-extracted stomatin assembles into two distinct architectures: a conical single-layer structure and a double-layer structure formed by two opposing cones (Figure S1). These two forms could not be clearly separated by a 10–50% glycerol gradient (Figure S1A). Data collection and image processing for cryo-EM yielded high-resolution reconstructions of both assemblies, hereafter termed STOM-single and STOM-double (Figure S2). STOM-single represented the majority of classified particles (∼80%) and was resolved at 2.2 Å (Figures 1A–D, S2, S3A–D), whereas STOM-double was less abundant but nonetheless resolved at 2.7 Å (Figures 1E–H, S2, S3E–H).

**Figure 1.**
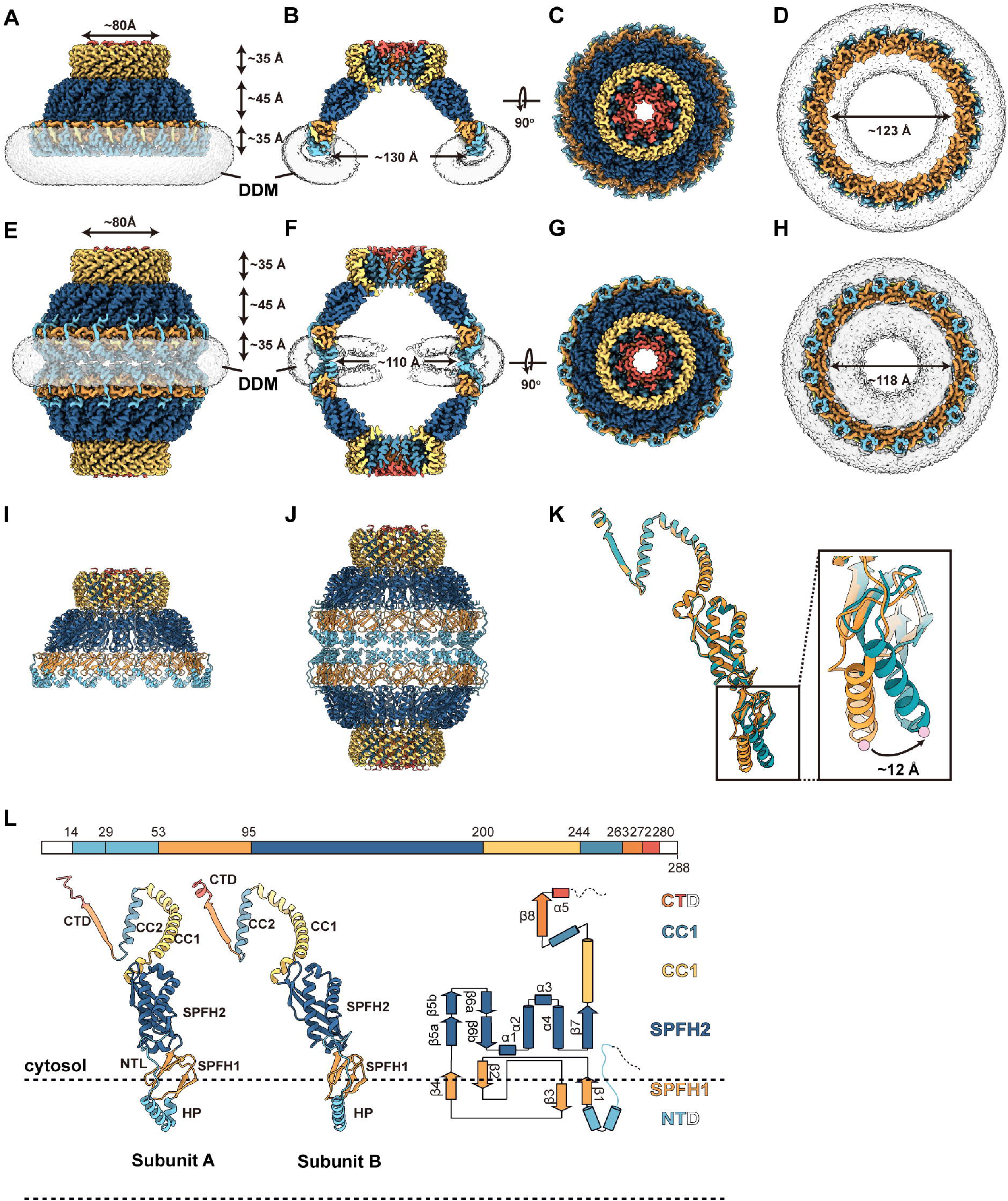
Overall structures of stomatin. **(A–D)** Cryo-EM density map of the STOM-single oligomer purified in DDM. The complex assembles into a bowl-shaped hexadecamer consisting of 16 subunits. The SPFH1 domain and intramembrane hairpin span ∼35 Å in height, the SPFH2 domain ∼45 Å, and the coiled-coil (CC) region ∼35 Å (A–B). The overall diameter of the hexadecamer is ∼130 Å (C), while the membrane area encircled by the SPFH1 domains measures ∼123 Å (D). **(E–H)** Cryo-EM density map of the STOM-double oligomer purified in DDM. The polymer consists of two bowl-shaped assemblies joined base-to-base via their N-terminal domains (NTDs). The height of each hexadecameric layer is consistent with that of STOM-single (E–F). The overall diameter of STOM-double is ∼110 Å (G), and the enclosed membrane area defined by the SPFH1 domains is ∼118 Å (H). **(I–J)** Atomic model of stomatin built on the cryo-EM density maps of STOM-single (I) and STOM-double (J). **(K)** Structural comparison of atomic models derived from the STOM-single (green) and STOM-double (orange) maps. The C-terminal regions align well, whereas a significant displacement is observed at the N-terminus, with a maximal shift of ∼12 Å in the hairpin domain. **(L)** Schematic representation of stomatin domain organization (top) and corresponding atomic model with topological diagram (bottom). NTD, N-terminal domain. HP, hairpin structure. NTL, N-terminal loop. SPFH1, SPFH1 domain. SPFH2, SPFH2 domain. CC1, coiled-coil domain 1. CC2, coiled-coil domain 2. CTD, C-terminal domain.

The cryo-EM map of STOM-single revealed a hexadecameric assembly comprising 16 stomatin subunits, each containing six domains: the N-terminal domain (NTD), SPFH1 and SPFH2 domains, coiled-coil domain 1 (CC1), coiled-coil domain 2 (CC2), and the C-terminal domain (CTD). The overall architecture resembles an inverted bowl anchored on the membrane, with a height of ∼105 Å and a bottom inner diameter of ∼130 Å (Figures 1A–B). By contrast, STOM-double consists of two single bowls joined base-to-base via their NTDs, resulting in opposite topologies between the two layers (Figure 1E). As only a few proteins have been reported to exhibit dual topologies in nature, such observations are more likely to represent artifacts of purification. But interestingly, STOM-double exhibits a smaller bowl diameter (∼110 Å) and an additional density corresponding to the N-terminal extramembrane region, which is absent in STOM-single (Figures 1E–F). Using the two maps, we built atomic models of stomatin (Figures 1I–J) and compared their conformations in the single- and double-layer states.

First, the two structures are highly consistent in the basal region of the bowl, which comprises a three-layer arrangement formed by the CC domain and CTD (Figure 2A–E). The outermost layer is composed of the CC1 domain, whose length varies among SPFH family members and may determine their oligomeric states. The middle layer contains the CC2 domain, and interactions between adjacent CC1 and CC2 domains stabilize the oligomeric architecture (Figures 2J–K). Proper angular positioning of CC1, SPFH1, and CC2 is essential for maintaining the tertiary structure and promoting subsequent oligomerization of STOM. Supporting this, targeted mutations at two pivotal proline residues located at the inter-domain interfaces (P200A and P245A) abolished STOM oligomerization (Figures 2J and 2M). The innermost layer forms a hydrophobic channel of ∼25 Å in diameter, constituted by β8-barrel of the CTD (Figures 2B and 2D). During data processing, we found that while STOM exhibits overall C16 symmetry, its CTD region displays only C8 symmetry, with β8 segments of adjacent subunits alternating between inward and outward orientations (Figures 2F–H). Notably, despite identical primary sequences, these subunits adopt distinct secondary structures: one forms an inward-facing α-helix that narrows the channel diameter to ∼17 Å, whereas the other forms an outward-projecting loop (Figures 2A, 2I and 2J). A similar pattern of alternating subunit conformations is observed in heteromeric SPFH family proteins, such as HflK/C^28,29^, erlin1/2, flotillin1/2^30^ and prohibitin1/2^31^. Remarkably, stomatin exhibits this property as a homo-oligomer, highlighting its two alternative conformations for its proper assembly.

**Figure 2.**
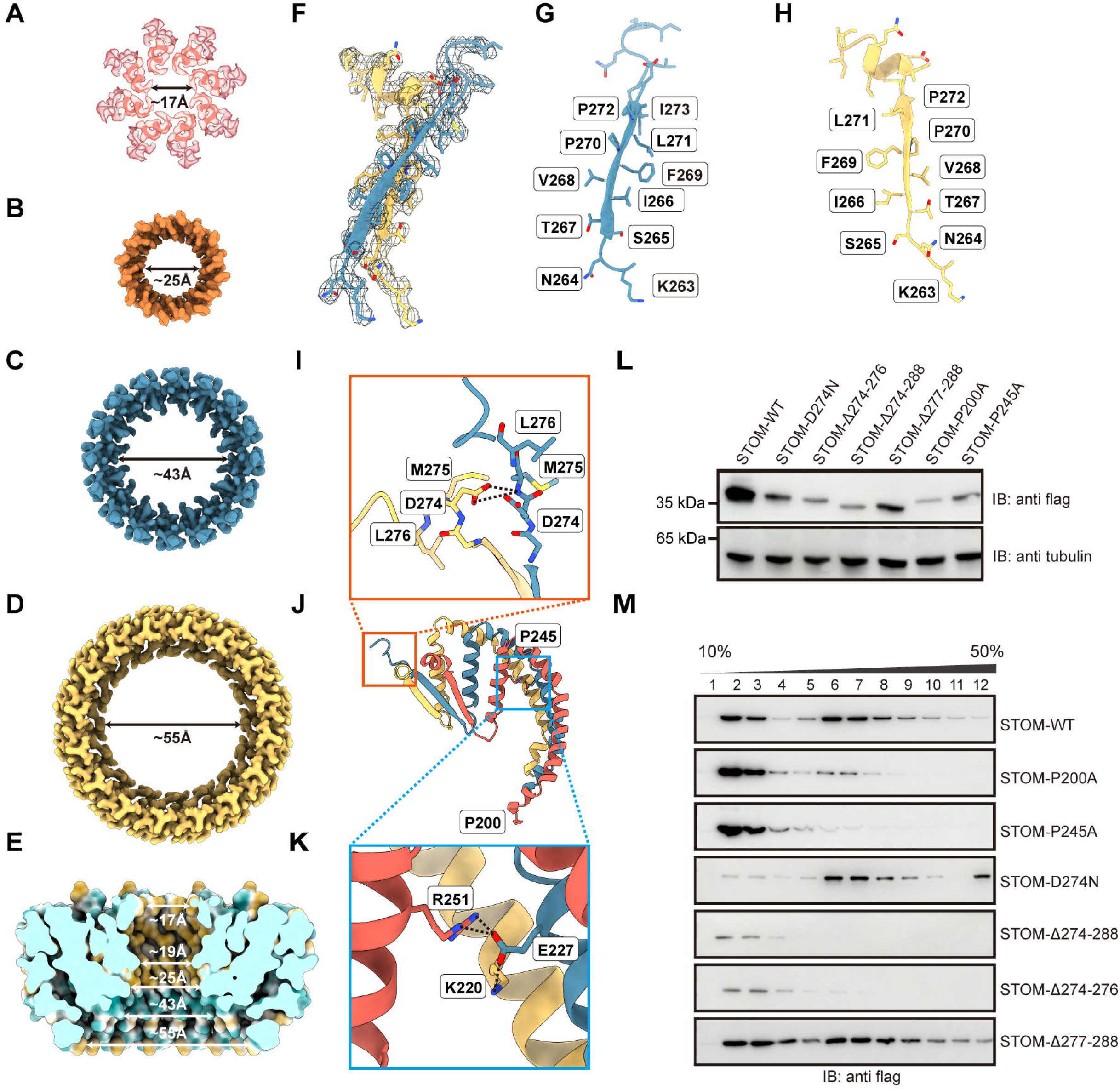
Structural features and mutational analysis of the basal region of the stomatin bowl. **(A–D)** Cryo-EM density maps of the bowl-bottom structure. From bottom to top: the outermost CC1 domain forms a layer with a diameter of ∼55 Å (D); the second layer consists of CC2, with a diameter of ∼43 Å (C); the innermost layer is formed by the β8-barrel of the CTD, with a diameter of ∼25 Å (B); and the lid structure beyond the β8-barrel contains a central hole with a diameter of ∼17 Å (A). **(E)** Hydrophilic and hydrophobic surface representation of the bowl-bottom region, revealing a central hydrophobic channel with a minimum diameter of ∼19 Å. **(F–H)** Cryo-EM density map (F) and atomic model (G–H) showing that adjacent β8-barrels alternate between inward- and outward-facing conformations. **(I–J)** Atomic model illustrating distinct secondary structures of the β8 segments: one subunit adopts an inward-facing α-helix, while the neighboring subunit forms an outward-projecting loop. **(K)** Structural model demonstrating that E227 in CC1 forms stabilizing salt bridges with acidic residues in CC2 and CC1 of adjacent subunits, thereby supporting oligomerization. **(L–M)** Mutational analyses of oligomerization. Protein expression was verified in cells (L), and 10%–50% glycerol gradient fractionation was used to examine the distribution of stomatin proteins (M). Stomatin-WT distributes into both high-molecular-weight multimers and low-molecular-weight monomers/oligomers. Two conserved proline residues (P200 and P245) are required for oligomerization, as their mutations (P200A, P245A) abolished polymer formation. Truncation of the CTD also impaired multimerization, with a strong effect observed at residues 274–276 (highlighted in I). D274N increased the proportion of multimers, consistent with reduced inter-subunit charge repulsion.

Previous studies have shown that the CTD (residues 264–288), particularly the β8-barrel region (264–273), is essential for stomatin oligomerization^23,32^. Our truncation experiments further revealed that the more distal C-terminal segment (274–276; DML) is also critical for oligomerization (Figure 2M). Structural analysis indicates that D274 forms a polar interaction with the backbone of M275 (Figure 2I) and the D274N mutation appeared to increase the proportion of polymers, likely by alleviating charge repulsion between D274 residues of adjacent subunits (Figures 2I and 2M).

### 2. Distinct conformations of stomatin define alternative modes of membrane insertion

In addition to the basal region of the stomatin bowl-like structure (CC1, CC2 and CTD), the SPFH2 domains of the STOM-single and STOM-double also remain highly consistent in conformation. The difference between these two assemblies primarily lie in the membrane-inserting region. Notably, substantial variation exists in the NTDs across SPFH family proteins. Whereas HflK/C^28,29^ and erlin1/2^33,34^ possess full transmembrane helices at their N-termini, the membrane insertion of stomatin primarily depends on its SPFH1 domain (Figures 3A–B), because the N-terminal sequence folds into an α-helical hairpin (HP), stabilized by a conserved proline residue (P47) located at the center of the helix (Figure 3C). This proline is strictly conserved among stomatin family members—including stomatin like protein 1 (STOML1), STOML3, and podocin (PODO)—indicating that these proteins likely adopt similar hairpin conformations (Figure S4A). In the STOM-double structure, the HPs of the upper and lower layers engage in hydrophobic interactions, thereby stabilizing the assembly into a D16-symmetric architecture (Figure 3E).

**Figure 3.**
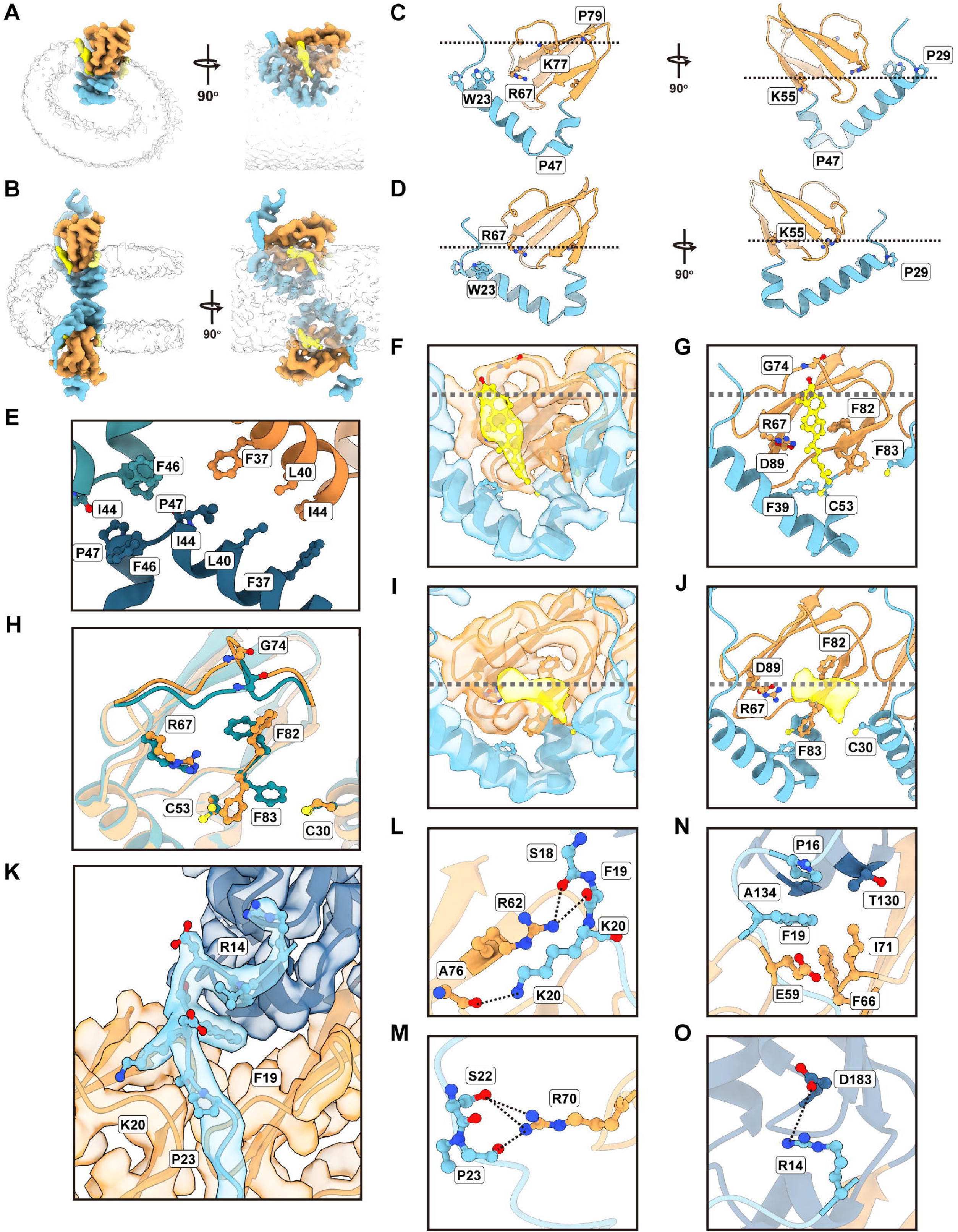
Membrane insertion and lipid interactions of stomatin. **(A–D)** Cryo-EM maps (A, B) and atomic models (C, D) of the intramembrane regions of STOM-single (A, C) and STOM-double (B, D). Positively charged residues that define the membrane insertion boundary are indicated in (C) and (D). DDM, white; NTD, light blue; SPFH1, orange; extra density, yellow. **(E)** Hydrophobic interactions between hairpins (HPs) of adjacent layers stabilize the D16-symmetric assembly of STOM-double. **(F–G)** The membrane-inserting region of STOM-single binds excess density (yellow) consistent with cholesterol (F), with surrounding amino acids shown in (G). **(H)** Comparison of the transmembrane regions of STOM-single and STOM-double reveals principal differences at F82 and F83, whose side chains adopt distinct deflections. **(I–J)** The membrane-inserting region of STOM-double also binds excess density (yellow, I), with surrounding amino acids shown in (J). **(K–O)** Interactions between the N-terminal loop (NTL) and SPFH domains in STOM-double. Clear electron density is observed for residues 14–20 (K). Stabilization of the NTL involves polar interactions of R62 and R70 with the NTL backbone (L, M), hydrophobic insertion of F19 (N), and a salt bridge between R14 and D183 (O).

Beyond the hairpin, the SPFH1 domain also embeds partially into the membrane (Figures 3A–B), a feature conserved with HflK/C^28,29^ and erlin1/2. Importantly, HP elements in stomatin do not interact directly with one another; instead, oligomerization is mediated by SPFH1–SPFH1 interactions, which partition the membrane into discrete functional domains. These interactions occur mainly through loops connecting β-barrels (Figure S4B) and are conserved across both STOM-single and STOM-double conformations (Figure S4C). Relative to STOM-single, STOM-double exhibits an inward deflection originating just below these loops in the SPFH1 domain. This deflection gradually increases, reaching ∼12 Å at the HP region (Figure 1K), thereby shrinking the circular membrane area organized by stomatin from about 120 nm^2^ to 110 nm^2^ (Figures 1D and 1H) and potentially deforming the surrounding membrane. Positively charged residues within both the HP and SPFH1 domains further define the membrane boundary. Interestingly, the two conformations embed differently into DDM micelles: STOM-single is inserted more deeply than STOM-double at the outer surface of the bowl (Figures 3C–D).

Three cysteines (C30, C53, C87) are present in the intramembrane region of stomatin and can undergo palmitoylation, enhancing membrane anchoring. Previous studies identified C30 as the primary palmitoylation site, with C87 also modified to a lesser extent^24^. Consistent with this, the STOM-single map revealed strong extra density adjacent to C30 and C87.

Importantly, we observed additional density on the surface of SPFH1 domain that closely resembled cholesterol (Figures 3F–G). The STOM-double structure revealed a strikingly different conformation (Figure 3H). Here, the extra density is in a different position, mimicking a head group of a lipid molecule (Figures 3I–J).

In the STOM-double structure, the N-terminal loop (NTL) preceding the helical HP extends upward to interact with the SPFH1 and SPFH2 domains of its own subunit, as well as the SPFH1 domain of the adjacent subunit (Figure 3K). Within SPFH1, R62 and R70 established polar interactions with the NTL backbone, stabilizing the otherwise flexible loop (Figures 3L–M). In addition, F19 in the NTL inserted into a hydrophobic pocket formed between SPFH1 and SPFH2 (Figure 3N), while R14 engaged in a salt bridge with D183 of SPFH2 (Figure 3O), further reinforcing the NTL binding.

Together, these conformational distinctions indicate that STOM-double is not generated merely by fusion of two STOM-single particles during purification. Instead, the two structures likely represent distinct functional states of stomatin oligomers.

### 3. Stomatin Organizes Cholesterol-enriched Membrane Microdomains through Self-Organization

To further investigate the behavior of stomatin on membranes, we reconstituted DDM-purified protein into liposomes composed of POPC, DOPS, and cholesterol (mass ratio 8:1:1), followed by cryo-sampling and data collection. Under these conditions, stomatin preferentially inserted into both leaflets of the lipid bilayer, forming characteristic double-layer structures (Figure S5-6). On smaller liposomes with high curvature, stomatin particles appeared mainly as isolated complexes, whereas larger liposomes with lower curvature supported the formation of clustered aggregates. Notably, stomatin diameter was largely insensitive to membrane curvature, as refinement of particles from liposomes of varying curvature yielded a uniform structure at 2.5 Å resolution (Figure S5). Comparison of liposome-reconstituted stomatin structures with ones in DDM micelles showed that the interfaces between adjacent subunits were all preserved. The membrane-insertion region of STOM-liposome more closely resembled STOM-double, although with slightly less contraction. Intriguingly, we detected no additional density in the membrane-insertion region, indicating that this pocket is affected by the conformation and orientation of the SPFH1 domains.

Interestingly, the circular membrane area encircled by stomatin oligomers appeared flattened (Figure S5B). Cryo-EM raw images and 2D classifications confirmed that stomatin assemblies could regularly cluster on liposome membranes, locally reducing membrane curvature (Figures 4A–B). We attempted to reconstruct the clustered assemblies but did not yield a high-resolution structure (Figure S6), likely due to variable spacing between adjacent stomatins. Collectively, these observations suggest that stomatin oligomers promote the formation and stabilization of larger membrane domains through self-organized clustering.

**Figure 4.**
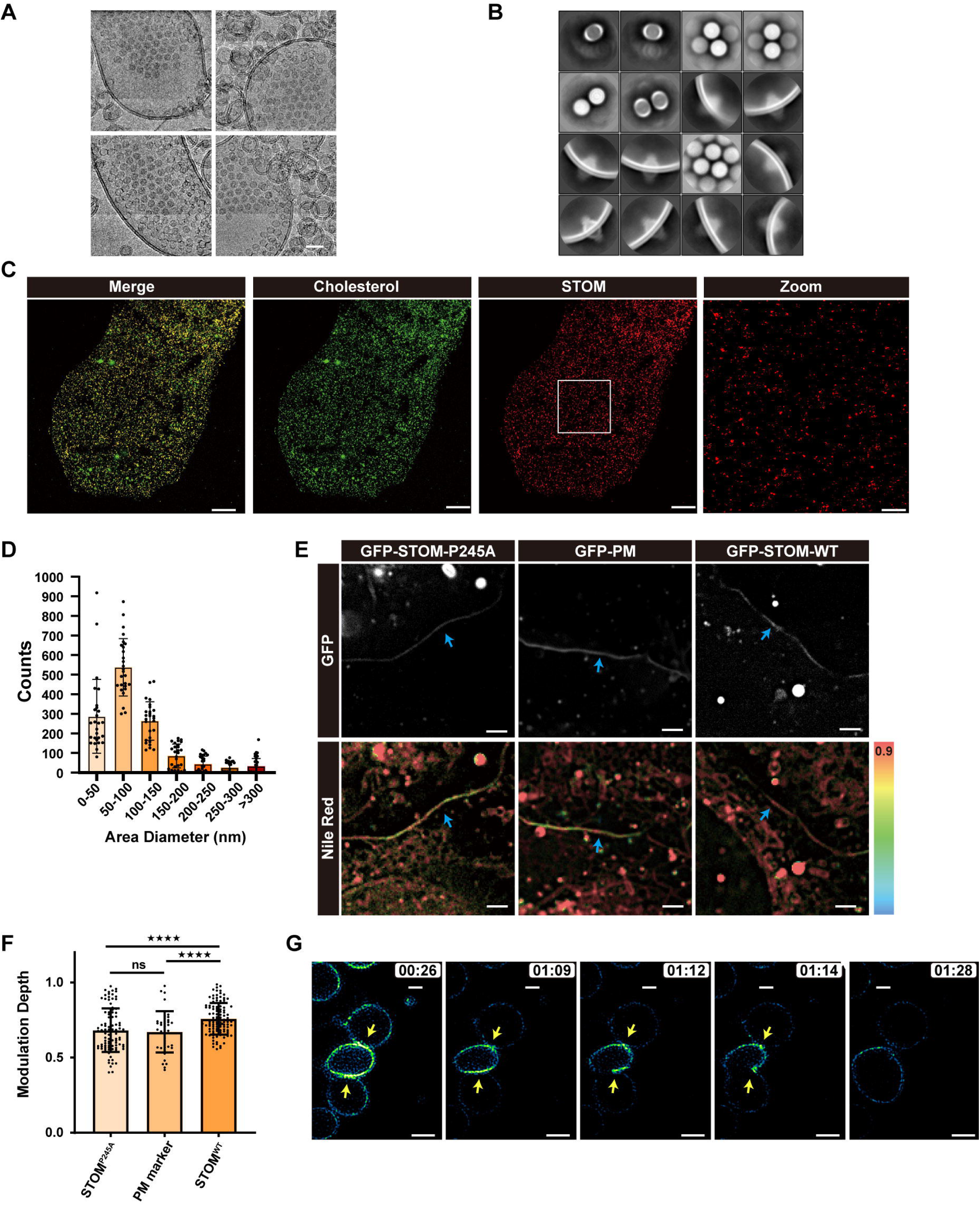
Stomatin clusters on membranes enhance lipid order and promote membrane remodeling. **(A)** Cryo-EM micrograph of stomatin reconstituted into POPC:DOPS:cholesterol (8:1:1) liposomes, showing the formation of stomatin clusters on the membrane. Scale bar, 50 nm. **(B)** STOM particles located in low-curvature membrane regions were selected for 2D classification (box size, 640 pixel). STOM particles appeared regularly arranged on the membrane. **(C–D)** U-2 OS cells were incubated with 25-NBD-cholesterol–BSA to replace cholesterol and subsequently immunostained for endogenous stomatin. Confocal microscopy revealed cholesterol enrichment at stomatin-positive sites (first three pictures of C; scale bar, 10 μm). STED super-resolution imaging confirmed cluster-like stomatin distribution (Zoom picture of C; scale bar, 2 μm). Cluster size and number were quantified in FIJI (D), showing diameters primarily within 50–150 nm, with some clusters >300 nm. Data represent three independent experiments (n = 27). **(E–F)** Polarization-resolved SIM analysis of U-2 OS cells expressing GFP-STOM. Representative images show GFP localization and Nile Red polarization mapping (E; scale bar, 2 μm). Blue arrows indicate analyzed membrane regions. Pseudo-color coding of Nile Red indicates lipid order: red, higher modulation depth (greater order); yellow, lower modulation depth (reduced order). Quantification of modulation depth (F) demonstrated significantly increased order in GFP-STOM-WT compared with GFP-PM (P < 0.0001) and GFP-STOM-P245A (P < 0.0001), whereas GFP-PM and GFP-STOM-P245A showed no significant difference (P = 0.6955). Data were obtained from three independent experiments; each point represents one membrane region. Statistical analysis was performed using an unpaired t-test (ns, not significant; ****, P < 0.0001). **(G)** Transient dehydration of A549 cells stably expressing GFP-STOM-WT was used to mimic stomatin-induced membrane remodeling. Removal of DMEM for 5 min followed by re-addition caused cell shrinkage, leaving GFP-STOM–enriched vesicles at the cell periphery. Adjacent vesicles rapidly fused with stomatin accumulation at the fusion sites (yellow arrows), accompanied by membrane fragmentation and reorganization at cluster edges.

To test whether such self-organization occurs in cells, we labeled live U-2 OS cells with the 25-NBD-cholesterol and performed immunofluorescence using anti-stomatin antibodies, followed by confocal and stimulated emission depletion (STED) microscopy of the basal plasma membrane. Stomatin was distributed as discrete cholesterol-enriched patches (Figure 4C), typically ranging from 50–150 nm in diameter, with some clusters exceeding 300 nm (Figure 4D).

Because highly ordered lipid domains exhibit more uniform Nile-Red orientation, they display stronger polarization responses and greater modulation depth. We therefore assessed lipid order using Spectrum and Polarization Optical Tomography (SPOT), a technique commercialized as the pGC modality in the polar-structured illumination microscopy (p-SIM) system^35^. In U2-OS cells transiently transfected with plasmids, we expressed GFP-tagged stomatin-WT (GFP-STOM-WT) or the polymerization-deficient P245A mutant (GFP-STOM-P245A), while GFP fused to a PM localization motif (GFP-PM) served as a negative control. The polarization response of Nile Red in membrane regions enriched with these fusion proteins was measured to evaluate lipid order. Regions enriched in GFP-STOM-WT aggregates exhibited significantly greater modulation depth than GFP-PM controls, indicative of increased lipid order. In contrast, GFP-STOM-P245A failed to enhance lipid order (Figures 4E–F). These results support a model in which stomatin polymerization drives the assembly of cholesterol-rich lipid raft microdomains.

Finally, we transiently dehydrated U2-OS cells stably expressing GFP-STOM-WT to simulate the membrane remodeling dynamics that stomatin-cluster may promote. Removal of DMEM for 5 min followed by readdition caused adherent cells to shrink, leaving GFP-STOM-WT–rich membrane vesicles at the periphery. These adjacent vesicles rapidly fused, with stomatin accumulation at fusion sites, accompanied by membrane fragmentation and reorganization at the stomatin-cluster edges (Figure 4G). This process was captured by continuous SIM imaging (Movie S1). Together, these findings suggest that stomatin-rich membrane regions are mechanically rigid, while their edges are more prone to lipid perturbations, providing a potential molecular mechanism by which stomatin promotes membrane fusion^13,14^ and remodeling.

### 4. Identification of Cargo Proteins Enriched in FMMs Organized by Stomatin

Function Membrane microdomains, typically lipid rafts, recruit and enrich raft-like proteins due to their unique lipid composition, thereby contributing to important cellular processes such as signal transduction. Although previous studies have identified several stomatin-interacting proteins potentially enriched in stomatin-organized FMMs^12,15,17,18^, systematic analyses have been lacking. To identify proteins enriched in FMMs organized by stomatin—hereafter referred to as STOM cargo proteins—we designed mass spectrometry (MS)–based approaches combining co-immunoprecipitation (co-IP) MS and enzymatic proximity labeling MS (Figures 5A–B).

**Figure 5.**
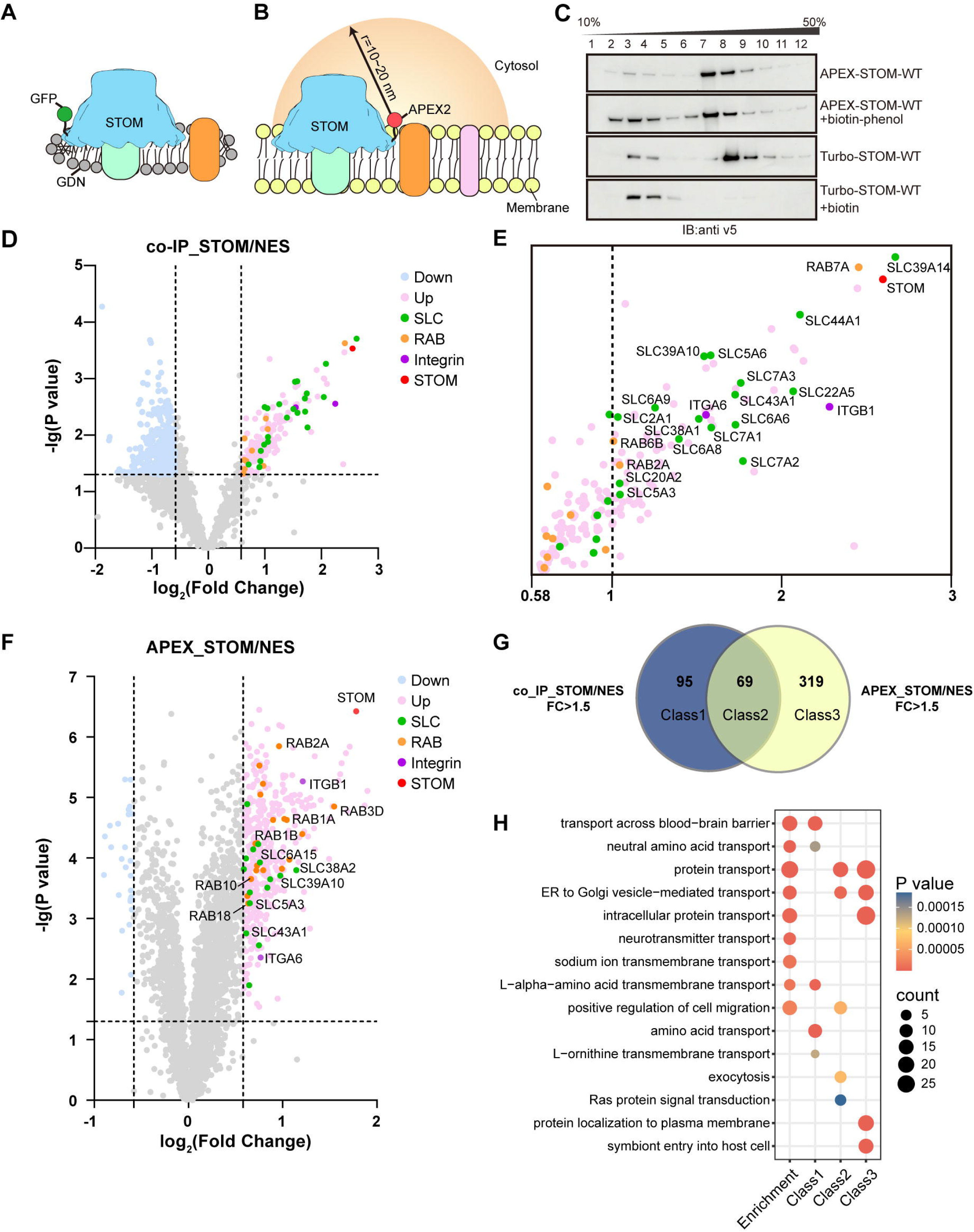
Proteomic profiling of STOM-associated cargo proteins by co-IP and proximity labeling mass spectrometry (MS). **(A–B)** Experimental design. GFP was fused to the N terminus of STOM for co-IP MS (A), and APEX2 was fused to the N terminus of STOM for proximity labeling MS (B). **(C)** Multimerization assessment of STOM fusion constructs. APEX2 did not significantly affect STOM multimerization, whereas TurboID largely abolished STOM multimers after 10 min of biotin labeling. **(D–F)** Volcano plots of MS results. Data represent three biological replicates. Proteins enriched by co-IP (D) or APEX2 labeling (F) were quantified with IBT-16plex reagents and analyzed by two-sample t-test in Perseus. The ordinate shows –log(P value), with the dotted line indicating 1.301 (P = 0.05). In co-IP (D), proteins highlighted in pink were >1.5-fold enriched in STOM-WT compared with NES; among these, SLC family proteins, RAB GTPases, and integrins were specifically labeled (E). In APEX2 (F), enriched proteins were defined as >1.5-fold, and overlapping SLCs, RABs, and integrins with co-IP enrichments were annotated with names. Enriched proteins are shown in pink; SLC family proteins, green; RAB GTPases, orange; integrins, purple. **(G)** Venn diagram showing overlap between co-IP and APEX2 enrichments. **(H)** Functional enrichment analysis of STOM-associated cargo proteins. Proteins enriched >1.5-fold in co-IP were labeled “enrichment,” and Class 1–3 groups were defined as in (F). Enrichment analysis was performed with DAVID, and the top-ranked biological pathways (BP) were selected for display.

For these experiments, we generated a series of stable HEK293T cell lines via lentiviral transduction. For the Co-IP MS analysis, we employed GFP-STOM-WT to capture stomatin and its interacting partners from cellular lysates. Additionally, we generated a cytoplasm-localized GFP-NES control by fusing GFP to a nuclear export signal (NES), enabling subtraction of GFP tag-derived contaminants. Enrichment was performed using a GFP nanobody. GDN was chosen for membrane extraction due to its ability to preserve the membrane region within the stomatin assemblies. This approach enriches cargo proteins interacting with stomatin directly or embedded in the membrane region within the stomatin bowl-like structure (Figure 5A).

We also applied proximity labeling MS to profile STOM cargo proteins (Figure 5B). The labeling enzyme was fused at the N-terminus of stomatin, such that labeled proteins should reside outside the closed stomatin bowl. Two enzymes, APEX2 and TurboID, were tested. APEX2 rapidly biotinylates exposed tyrosine residues within ∼10–20 nm of its active site in ∼1 minute^36,37^,whereas TurboID biotinylates lysine residues within ∼5–10 nm but requires ∼10 minutes for effective labeling^36,38^. Notably, TurboID labeling induced depolymerization of STOM assemblies, whereas APEX2 labeling had neglectable effect (Figure 5C), likely due to shorter labeling times and the smaller steric interference at the interface between adject subunits. Consequently, we employed APEX2 for subsequent proximity labeling MS experiments.

We first performed enrichment analysis of proteins showing more than 1.5-fold enrichment in GFP-STOM versus GFP-NES co-IP MS, and the most significantly enriched group was the solute carrier (SLC) family transporters (Figures 5D, 5E and 5H). Furthermore, we compared the enriched proteins (fold change > 1.5) of identified by co-IP and proximity labeling MS, and found that their union (class2)—likely representing cargo proteins residing outside the stomatin bowl—showed strong enrichment of vesicular transport proteins, particularly those mediating trafficking between the endoplasmic reticulum (ER), Golgi apparatus, and late endosomes (Figures 5F–H). This set included multiple RAB family GTPases, suggesting a role for STOM in intracellular transport (Figure 5F). Among these RABs, the Rab7a, which is the marker protein of late-endosome has been significantly enriched by stomatin (Figure 5D), consistent with previous reports showing that stomatin also localized on late endosomes^39–41^. Notably, we also detected abundant interactions with migration-related proteins, including integrins ITGB1 and ITGA6, at the outer surface of STOM assemblies (Figures 5D–H), suggesting a role for stomatin in cell migration.

Together, our MS analyses indicate that stomatin plays an important role in vesicle transport, solute channel regulation, and cell migration.

## Discussion

In this study, we determined the 16-mer structure of stomatin and identified a potential cholesterol-binding pocket within its membrane-inserting region. Using liposome reconstitution, we demonstrated that stomatin can self-assemble into organized arrays on membranes. Super-resolution imaging further revealed that stomatin organizes membrane microdomains of ∼50–150 nm or larger in diameter, which exhibit enhanced lipid order, providing direct experimental evidence that stomatin acts as a scaffolding protein for FMMs, which are enriched in saturated lipids. Mass spectrometry analyses demonstrated that stomatin interacts with a range of cargo proteins, such as SLC transporters, RAB GTPases, and integrins, suggesting that it participates in diverse cellular processes. Based on these structural and biochemical findings, together with previous studies, we propose the following molecular model for stomatin function.

When reconstituted on artificial membrane liposomes, stomatin spontaneously assembled into clusters, suggesting a similar role in organizing FMMs in cells. Such clustering enhances lipid order in the local membrane region, thereby generating a distinct lipid environment that may influence the conformation and activity of raft-associated proteins. In addition, clustering increases the local concentration of stomatin cargo proteins, potentially facilitating protein concentration–dependent physiological processes. For example, co-IP and proximity labeling MS together confirmed that stomatin interacts with integrins, which play a central role in cell migration^42,43^. Stomatin may promote integrin clustering, with important implications for the regulation of cell migration.

Focusing on the stomatin oligomer, we propose that its largely closed bowl-like structure acts as a barrier that can trap certain proteins and regulate their activity. Mass spectrometry results indicate that the most abundant of these proteins are SLC family transporters. Consistent with this, previous studies have shown that stomatin inhibits the function of GLUT1^16,17^, a glucose transporter in this family. Stomatin also suppresses the ion transport activity of the gap junction (GJ) protein Panx1^12^; while recent study identified that stomatin can restrain some of GJ channels within the bowl in primary *Caenorhabditis elegans* cells by cryo-electron tomography (cryo-ET)^44^. Our structural analysis reveals that the stomatin polymer forms a nearly closed assembly, leaving only a hydrophobic pore form by β-barrels at the C-terminus (∼19 Å), which may be further occluded by the following C-terminal segments. We therefore speculate that stomatin may regulate channel activity by affecting its substrate permeability or altering its membrane environment. Certainly, this barrier function may also operate by shielding cargo proteins from external interactions, thereby limiting specific cellular processes.

Together, these findings position stomatin as a dynamic organizer of functional membrane microdomains, participating in essential cellular processes such as solute transport, vesicle trafficking, and cell migration by regulating its cargo proteins.

## EXPERIMENTAL MODEL AND STUDY PARTICIPANT DETAILS

### Cell culture and treatments

HEK293T (CRL-3216) were obtained from ATCC. U2os cells were generously provided by P. Xi (Peking University, Beijing). All cells are maintained in DMEM (Gibco) supplemented with 10% FBS and 1% penicillin/streptomycin at 37°C incubator with 5% CO2. HEK293F (Thermo Fisher Scientific, A14528) cells were cultured in SMM 293-TII (Sino Biological), in a 37□°C and 5% CO2 incubator shaking at 120 rpm.

## METHOD DETAILS

### Cloning, expression and purification of the stomatin complex

The genes of stomatin were amplified from the cDNA library of HEK293F cells. The stomatin gene was fused with a Twin-Strep tag and cloned into pcDNA3.1 vector. Plasmid DNA and PEI (Polysciences) were transfected into HEK293F cells at a density of approximately 1.5×10□ cells/mL, using a mass ratio of 1:3 (plasmid:PEI). Transfection was performed with 1 mg of plasmid DNA per liter of culture.

Cells were harvested by centrifugation and washed once with PBS. The cell pellet was resuspended in lysis buffer (40 mM HEPES-KOH, pH 7.4, 150 mM NaCl, 10% glycerol) supplemented with 1× cocktail protease inhibitor (Mei5bio) and lysed by sonication. The lysate was centrifuged at 1,300 × *g* for 20 min to remove cell debris, and the supernatant was centrifuged at 45,000 rpm for 1 hr in a Type 70Ti rotor (Beckman Coulter) to collect the membrane fraction. The pelleted membrane fraction was solubilized in lysis buffer containing 1% (w/v) n-dodecyl-β-d-maltoside (DDM, Anatrace) at 4°C for 4 hr. The supernatant was collected by centrifugation at 45,000 rpm for 30 min in the Type 70Ti rotor (Beckman Coulter). The supernatant was incubated with Strep-Tactin 4Flow (iba) at 4°C for 2 hr with rotation, followed by washing with 100 c.v. of wash buffer (lysis buffer supplemented with 1× cocktail protease inhibitor and 0.0174% (w/v) DDM) and elution with wash buffer containing 20 mM d-Desthiobiotin (Sigma-Aldrich). The eluate was concentrated with a 100-kDa cutoff spin concentrator (Merck Millipore) and loaded onto a 10%–50% glycerol density gradient and centrifuged at 100,000 × *g* for 14 hr in a SW41 Ti rotor (Beckman Coulter). All fractions of the gradient were collected and analyzed by SDS-PAGE. The peak fraction was concentrated for cryo-EM sample preparation.

### Negative staining electron microscopy examination

4 uL of sample at a concentration of approximately 0.2 mg/mL was loaded onto a copper grid and stained with 2% uranyl acetate. The grid was examined using a FEI Tecnai G2 Spirit at 120 kV.

### Cryo-EM sample preparation and data collection

4 μL of sample with a concentration of approximately 10 mg/mL was loaded onto a glow-discharged gold grid (Quantifoil, R2/1, 300 mesh), waited for 4 s, and blotted for 1□s with –1 blot force using an FEI Vitrobot at 6□°C and 100% humidity.

Data were collected using a FEI Titan Krios (with Gatan K2 summit camera). Movies were collected using SerialEM^45^ at 130,000× magnification (pixel size of 1.052□Å) with the defocus range varying from −0.7 μm to −1.3 μm. Each movie contained 32 frames with a dose rate of 8 e^−^/Å^2^/s for a total exposure time of 8□s.

### Cryo-EM data processing

The motion correction and electron-dose weighting of the movies were performed with MotionCor2^46^. The CTF parameters were estimated with the program of CTFfind4.1^47^. Particle picking, extraction, classification and refinement were done with RELION-5.0^48^.

The particles selected by the Laplace-Gaussian algorithm were subjected to multiple rounds of 2D classification and 3D classification. The good classes were selected for 3D refinement with C16 symmetry. For the calculation of the C-terminus with poor resolution, the center of the particle is moved to the C-terminus, 3D classification (skip alignment with C8 symmetry) is performed, and several good classes are selected for mask-based 3D refinement.

### Model building

The atomic model of stomatin was built *de novo* in Coot^49^ based on the cryo-EM density map. The construction of the C-terminus model uses the local map generated by particle center shifting. Model refinement was done using real-space refinement in Phenix^50^. The figures preparation were performed with UCSF Chimera^51^ and ChimeraX^52^.

### Reconstitution of stomatin into liposomes

Liposome preparation were performed as previously described^53^. POPC, DOPS, and cholesterol (8:1:1, mass ratio) were dissolved in chloroform and dried in a rotary vacuum evaporator at 38 °C water bath for 6 h to remove residual solvent. The dried lipid film was resuspended in reconstitution buffer (25 mM Na-PIPES, pH 7.2, 150 mM NaCl) to a final concentration of 40 mg/mL. The suspension was sonicated for 10 min and subjected to eight freeze–thaw cycles (liquid nitrogen-water bath at 40□°C) to reduce multilamellar vesicle formation. Liposomes were subsequently extruded through Nuclepore filters (100 nm pore size) using an Avanti Mini-Extruder to obtain liposomes of ∼100 nm in diameter.

Blank liposomes (3.2 mg) were solubilized with 0.5% n-dodecyl-β-D-maltoside (DDM) at room temperature for 1 h, followed by incubation with purified stomatin protein (0.8 mg) at 4 °C for 1 h. The protein–lipid–detergent mixture was applied to a gravity column packed with 18□ml Sephadex G-50 resin (superfine, Sigma-Aldrich) for detergent removal. Proteoliposomes eluted between 4.5 and 7.0 mL and were concentrated to <250 μL using a 100-kDa cutoff concentrator. The sample was adjusted to 20% iodixanol (Opti-Prep) and loaded onto a discontinuous iodixanol gradient (0%, 1%, 3%, 5%, 15%, 20%, 30%; top to bottom). After centrifugation at 50,000 × g for 16 h, empty liposomes were recovered in the 3–5% fraction, whereas reconstituted proteoliposomes were enriched in the 15–20% fraction. For subsequent cryo-EM sample preparation, the 15–20% fraction was collected and iodixanol was removed by seven cycles of dilution (1:5) and reconcentration. The sample was finally concentrated to ∼12 μL using a 100-kDa concentrator.

Grids were prepared using a Vitrobot Mark IV operated at 6 °C and 100% humidity. Quantifoil Au R 2/1 300-mesh holey gold grids were glow-discharged for 40 s. Four microliters of freshly prepared proteoliposomes were applied to the grid and incubated for 2 min before blotting. This procedure was repeated three times, with the final application performed after mixing the sample with 0.65 mM fos-choline-8 (Hampton Research). Grids were blotted for 5 s with blot force 0 and plunge-frozen into liquid ethane cooled by liquid nitrogen.

Data were collected on a Titan Krios-I cryo-electron microscope (Thermo Fisher Scientific) operated at 300 kV, equipped with a Gatan K3 direct electron detector and an energy filter, using EPU software. Images were acquired at 105,000× nominal magnification, corresponding to a pixel size of 0.85 Å. The defocus range was set to −1.5 μm. Each movie was recorded over 2.89 s with 40 frames, at a total electron dose of ∼60 e⁻/Å², using a slit width of 10 eV.

### Endogenous cholesterol labeling using 25-NBD-cholesterol

25-NBD-cholesterol powder was first dissolved in chloroform:methanol (3:1, v/v) to prepare a 100 nM stock solution. A 100 μL aliquot of the stock solution was evaporated under a gentle argon stream to form a thin lipid film. The dried film was dissolved in 50 μL ethanol, and 20 μL of this ethanol solution was added to 1 mL HDMEM (DMEM supplemented with 10 mM HEPES, pH 7.2) containing 100 μM BSA. After thorough mixing, the solution was diluted into 9 mL of HDMEM to generate the final BSA–cholesterol complex dye. The dye was freshly prepared and used immediately.

U-2 OS cells were cultured on 29 mm glass-bottom dishes (Cellvis, D29-20-1.5H) for treatment and immunofluorescence staining. Cells were plated 24 h prior to staining at ∼30% confluency. At the time of staining, cells reached ∼50% confluency. Freshly prepared 25-NBD-cholesterol dye was added to the cells and incubated for 30 min at 37 °C. After incubation, cells were washed three times with pre-warmed PBS (37°C) and fixed with 4% paraformaldehyde (PFA) for 15 min at room temperature, followed by three additional PBS washes. The cells were subsequently processed for immunostaining.

### Immunofluorescence staining and STED imaging analysis

The cell membrane was then permeabilized with 0.1% Triton X-100. Non-specific binding sites were blocked by incubating cells with 4% normal BSA in PBS for 1 hr at room temperature. Primary antibody incubation was performed overnight at 4°C using anti-stomatin polyclonal antibody (Proteintech) diluted in blocking solution. After three washes with PBS, cells were incubated with Abberior STAR RED-conjugated goat anti-rabbit IgG (H+L) cross-adsorbed secondary antibody (Abberior) for 40 min at room temperature, followed by three additional PBS washes.

Fluorescence images were acquired using a Leica TCS SP8 STED 3X microscope. For statistical analysis, images were uniformly acquired at a size of 23.25 μm × 23.25 μm with a pixel resolution of 22.73 nm × 22.73 nm. The collected images were processed using FIJI software^54^, where the diameter and number of puncta structures were quantified. Statistical graphs were generated using GraphPad Prism 10.

### Spectrum and Polarization Optical Tomography (SPOT)

Spectrum and Polarization Optical Tomography (SPOT) were performed as previously described^35^. To determine suitable constructs for imaging, GFP was fused to either the N-terminus of stomatin. The polymerization of the fusion proteins was retained normal. U-2 OS Cells were transfected using Lipofectamine 3000 (Thermo Fisher Scientific). The culture medium was replaced 8 h after transfection; and 30h after transfection, 1 µg/ml Nile Red was added into the culture medium 1 h before imaging. Fluorescence signals of GFP were collected under the 488-nm channel, while a 561 nm laser is used to excite NileRed with polar-Grid confocal (pGC) modality. The plasma membrane was distinguished based on its morphological features, and regions exhibiting both GFP and Nile Red signals (membrane areas expressing the recombinant proteins) were selected. Curves were manually drawn on the intensity images, and modulation depth were quantified using FIJI software^54^. Statistical analyses were performed to compare membrane order across different regions, generated using GraphPad Prism 10.

### Mass Spectrometry Construction of stable cell lines

For the proximity labeling MS, APEX2/TurboID sequence was fused to the N-terminus of stomatin, and a V5 tag was placed at the C-terminus; For the co-IP MS, GFP sequence was fused to the N-terminus of stomatin. Recombinant constructs expressing stomatin were generated using a doxycycline-inducible Tet-On vector. A NES sequence fused with APEX2/ TurboID /GFP was used as a negative control.

Stable cell lines were established using a second-generation lentiviral system. Briefly, HEK293T cells (30% confluency) were transfected in 6-well plates with plasmids (pTRZR:VSVG:PAX2 = 2:1:2, total 2.5 μg DNA) and PEI transfection reagent (7.5 μg) in serum-free, antibiotic-free DMEM. After 12 h, the medium was replaced with DMEM containing 10% FBS, and viral supernatants were harvested after 48 h. Viral particles were clarified by centrifugation (1,300 × g, 5 min) and filtration (0.45 μm), mixed 1:1 with fresh DMEM supplemented with 10% FBS, and supplemented with polybrene (10 μg/mL). Target HEK293T cells were infected with viral preparations, and uninfected cells were maintained as controls. After 24 h, medium was replaced with fresh DMEM, and after an additional 12 h, puromycin selection (1 μg/mL) was initiated. Following complete death of control cells (typically 48 h), resistant cells were maintained with reduced puromycin concentration (0.5 μg/mL) for 2–3 passages to establish stable cell lines. Aliquots were frozen for long-term storage, and remaining cells were used for proximity labeling experiments.

### APEX2 labeling for mass spectrometry

Stable cell lines were seeded in 10-cm dishes and induced with doxorubicin (0.1 μg/mL) for 36 h. At >90% confluence, the medium was replaced with 5 mL pre-warmed complete medium containing 500 μM biotin-phenol (BP; from 500 mM DMSO stock). Cells were incubated at 37 °C for 30 min, followed by the addition of 5 μL 1 M H_2_O_2_ (final concentration 1 mM). The dishes were gently shaken to distribute H_2_O_2_, and labeling was allowed for 1 min. The reaction was quenched twice for 2 min each with freshly prepared quenching buffer (10 mM sodium ascorbate, 5 mM sodium azide, 5 mM Trolox in PBS; alternatively, 50 mM sodium ascorbate without azide).

### Co-immunoprecipitation for mass spectrometry

Stable GFP-stomatin and GFP-NES cell lines were used for co-IP MS analysis. Cells were seeded in 10-cm dishes and induced with doxorubicin (0.1 μg/mL) for 36 h. Membrane proteins were extracted from equal cell numbers using 1% (w/v) GDN. The supernatant was collected by ultracentrifugation at 186,000 × g for 10 min using a TLA-55 rotor (Beckman Coulter). Equal volumes of supernatant were incubated with NHS resin conjugated to GFP nanobody. Bound proteins were directly denatured in 4× SDS loading buffer at 37 °C for 30 min. After SDS-PAGE, gel pieces containing the proteins were excised to remove detergent interference prior to mass spectrometry analysis.

### Mass spectrometry sample preparation APEX2-based proximity labeling

Cells were washed twice with phosphate-buffered saline (PBS) following completion of the labeling reaction. Subsequently, cells were resuspended in 4% (w/v) sodium dodecyl sulfate (SDS) aqueous solution and lysed by sonication on ice. The resulting lysate was centrifuged at 15,000 × g for 10 min at 4 °C. To remove residual probes, proteins were precipitated by incubation in cold methanol at −80 °C overnight. The precipitated proteins were collected by centrifugation at 3,000 × g for 10 min at 4 °C, washed twice with cold methanol (−80 °C), and then redissolved in 800 μL of 0.5% (w/v) SDS aqueous solution.

Protein concentration was determined using the Pierce™ BCA Protein Assay Kit (Thermo, 23227). For enrichment, 50 μL streptavidin agarose resin (Thermo, 20347) was equilibrated with 1 mL PBS and then incubated with the protein solution at 25 °C for 3 h under gentle rotation. Beads were washed once with 1 mL of 0.5% SDS in PBS for 10 min with gentle rotation, followed by six successive washes with 1 mL PBS. After discarding the supernatant, the beads were resuspended in 500 μL of 6 M urea in PBS. Subsequently, 25 μL of 200 mM dithiothreitol (Sigma, D9163-5G) was added and incubated at 37 °C for 60 min. The mixture was cooled to room temperature, followed by the addition of 25 μL of 400 mM iodoacetamide (Sigma, I6125-5G) and incubation at 30 °C for 30 min in the dark.

The beads were then washed four times with 1 mL of 100 mM triethylammonium bicarbonate (Sigma, T7408-100 mL) and resuspended in 200 μL of triethylammonium bicarbonate buffer. Proteins were digested on-bead using 1 μg sequencing-grade trypsin (Promega, V5111) at 37 °C for 16 h with agitation at 1,200 rpm. Digested peptides were collected by centrifugation at 15,000 × g for 10 min and dried by vacuum centrifugation.

### Liquid chromatography-tandem mass spectrometry

Peptides were separated using a loading column (100 µm × 2 cm) and a C18 separating capillary column (75 µm × 15 cm) packed in-house with 1.9 μm C18 bulk packing material (Dr. Maisch GmbH, Germany). The mobile phases (A: water with 0.1% formic acid and B: 80% acetonitrile with 0.1% formic acid) were driven and controlled by a Dionex Ultimate 3000 RPLC nano system (Thermo Fisher Scientific). The LC gradient was held at 2% for the first 8 minutes of the analysis, followed by an increase from 2% to 8% B from 8 to 9 minutes, an increase from 8% to 25% B from 9 to 90 minutes, an increase from 25% to 40% B from 90 to 123 minutes, and an increase from 44% to 99% B from 123 to 128 minutes.

For the samples analyzed by Orbitrap Fusion LUMOS Tribrid Mass Spectrometer, the precursors were ionized using an EASY-Spray ionization source (Thermo Fisher Scientific) source held at +2.2 kV compared to ground, and the inlet capillary temperature was held at 320°C. Survey scans of peptide precursors were collected in the Orbitrap from 350-1600 Th, a maximum injection time of 50 ms, RF lens at 30%, and a resolution of 120,000 at 200 m/z. Monoisotopic precursor selection was enabled for peptide isotopic distributions, precursors of z = 2-7 were selected for data-dependent MS/MS scans for 3 seconds of cycle time, and dynamic exclusion was set to 15 seconds with a ±10 ppm window set around the precursor monoisotopic.

In HCD scans, an automated scan range determination was enabled. An isolation window of 0.7 Th was used to select precursor ions with the quadrupole. Product ions were collected in the Orbitrap with the first mass of 100 Th, a maximum injection time of 118 ms, HCD collision energy at 38%, and a resolution of 60,000.

### Data analysis

The MS data of the samples were analyzed with Thermo Proteome Discoverer 2.4 software. For protein identification, MS/MS spectra were searched against UP000005640 proteome reviewed database from Uniprot (20964 human proteins in total) and GFP (green fluorescent protein). Trypsin was used for digestion, and two trypsin missed cleavage sites were allowed. Carbamidomethylation at cysteine (+57.021 Da) and iBT 16-plex at lysine/N-terminal (+277.188 Da) were set as fixed modifications. Oxidation at methionine (+15.995 Da) and acetylation of N-terminal (+42.011 Da) were set as variable modifications. The false discovery rate (FDR) for peptide identification was calculated using the Percolator algorithm in the Proteome Discoverer workflow based on the search results against a decoy database and was set at 1% FDR. Protein quantification was calculated based on the relative intensities of reporter ions (114 (114.1277 Da), 115N (115.1248 Da), 115C (115.1311 Da), 116N (116.1281 Da), 116C (116.1344 Da), 117N (117.1315 Da), 117C (117.1378 Da), 118N (118.1348 Da), 118C (118.1411 Da), 119N (119.1382 Da), 119C (119.1445 Da), 120N (120.1415 Da), 120C (120.1479 Da), 121N (121.1449 Da), 121C (121.1512 Da), 122 (122.1482 Da)) extracted from tandem mass spectra. Proteins with unique peptides <2 or any reporter ion intensity “0” or score “0” were removed.

Enrichment fold were calculated by division using their normalized MS intensity. Normalization of APEX-labeled samples employed median protein intensity from respective channels, whereas GFP tag intensity was utilized for normalization of Co-IP samples.

P values were calculated by Perseus v1.5.5.3 software.

## Acknowledgements

We thank the Core Facilities at the School of Life Sciences, Peking University (PKU) for help with negative staining EM; the PKU Cryo-EM Plat form and J. Wang., J. Zhao. and B. Xu. in PKU Institute of Advanced Agricultural Sciences for cryo-EM data collection; the PKU High performance Computing Platform for help with computation; the National Center for Protein Sciences (PKU Branch) for assistance with STED imaging; the PKU Analytical Instrumentation Center for assistance with MS sample identification and Ms. W. Zhou for help with MS results analysis. We thank Airy Technologies Co. Ltd. for assistance with Polar-SIM imaging. This work was supported by National Natural Science Foundation of China (92354306 to N.G.).

## Author contributions

L.Y. purified stomatin and determined its cryo-EM structure. N.G. and L.Y. performed model building. L.Y. and C.W. reconstituted stomatin into liposomes under the guidance of B.X. M.L. carried out SPOT data processing under the guidance of P.X. L.Y. and X.Z. prepared MS samples, and X.Z. performed mass spectrometry identification. L.Y., X.Z., and N.G. wrote the manuscript.

N.G. and P.Z. supervised the study.

## Supplementary Materials

Figures. S1 to S6

Movie S1

Table S1

