## Supplement Fig & Tab for "Structural Role of Stomatin in Organizing Functional Membrane Microdomains"

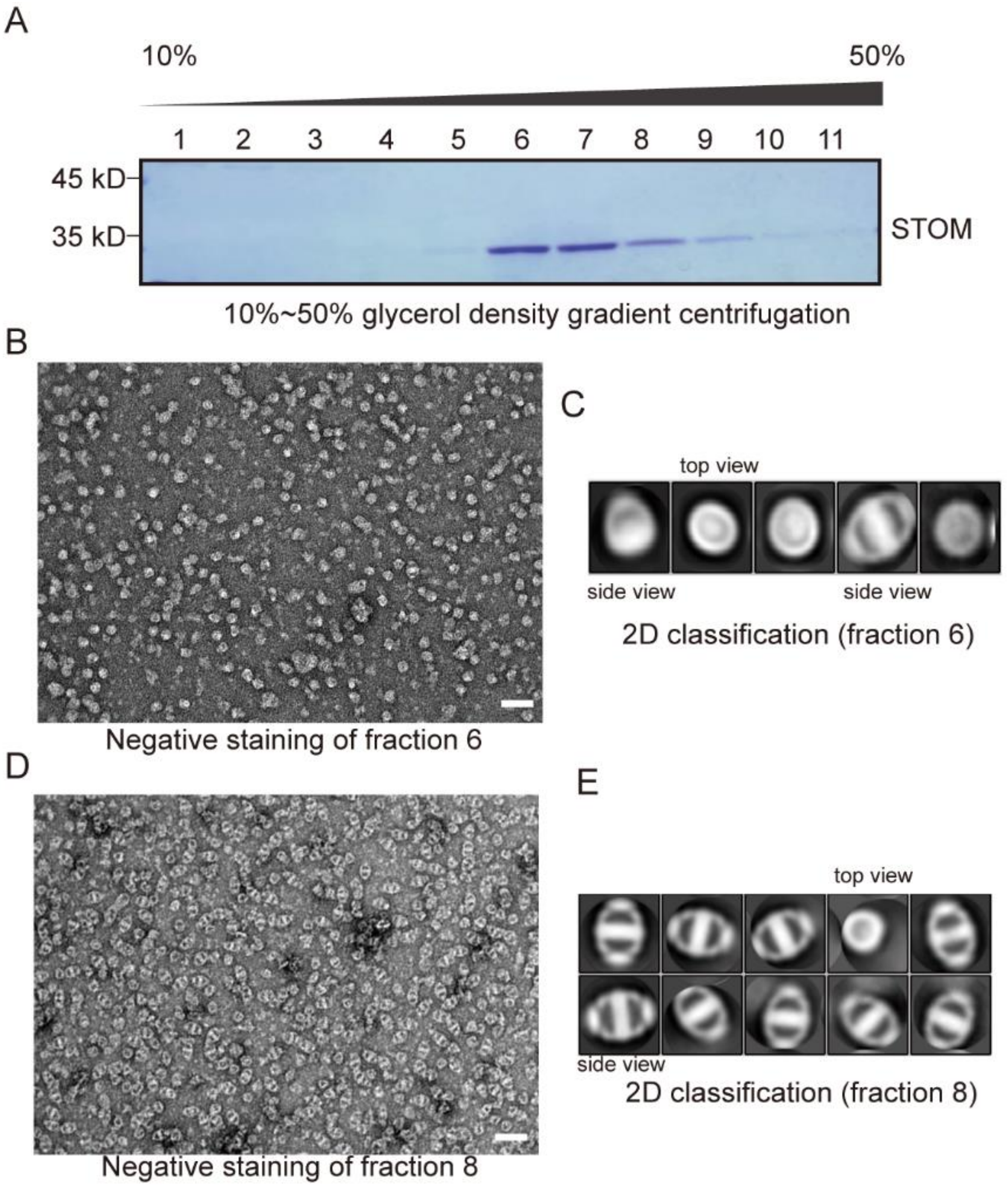

**Figure S1. Protein purification.**

**(A)** Eleven fractions collected from top to bottom after 10%–50% glycerol density gradient separation. The highest target protein content was detected in fractions 6–8, corresponding to ~33%–37% glycerol.

**(B–C)** Negative-stain transmission electron microscopy (TEM) of purified stomatin protein from fraction 6. Representative micrographs (B) and 2D classification results (C) are shown. A total of 100 micrographs were collected. Scale bar, 50 nm.

**(D–E)** Negative-stain TEM of stomatin protein from fraction 8. Representative micrographs (D) and 2D classification results (E) are shown. A total of 100 micrographs were collected. Scale bar, 50 nm.

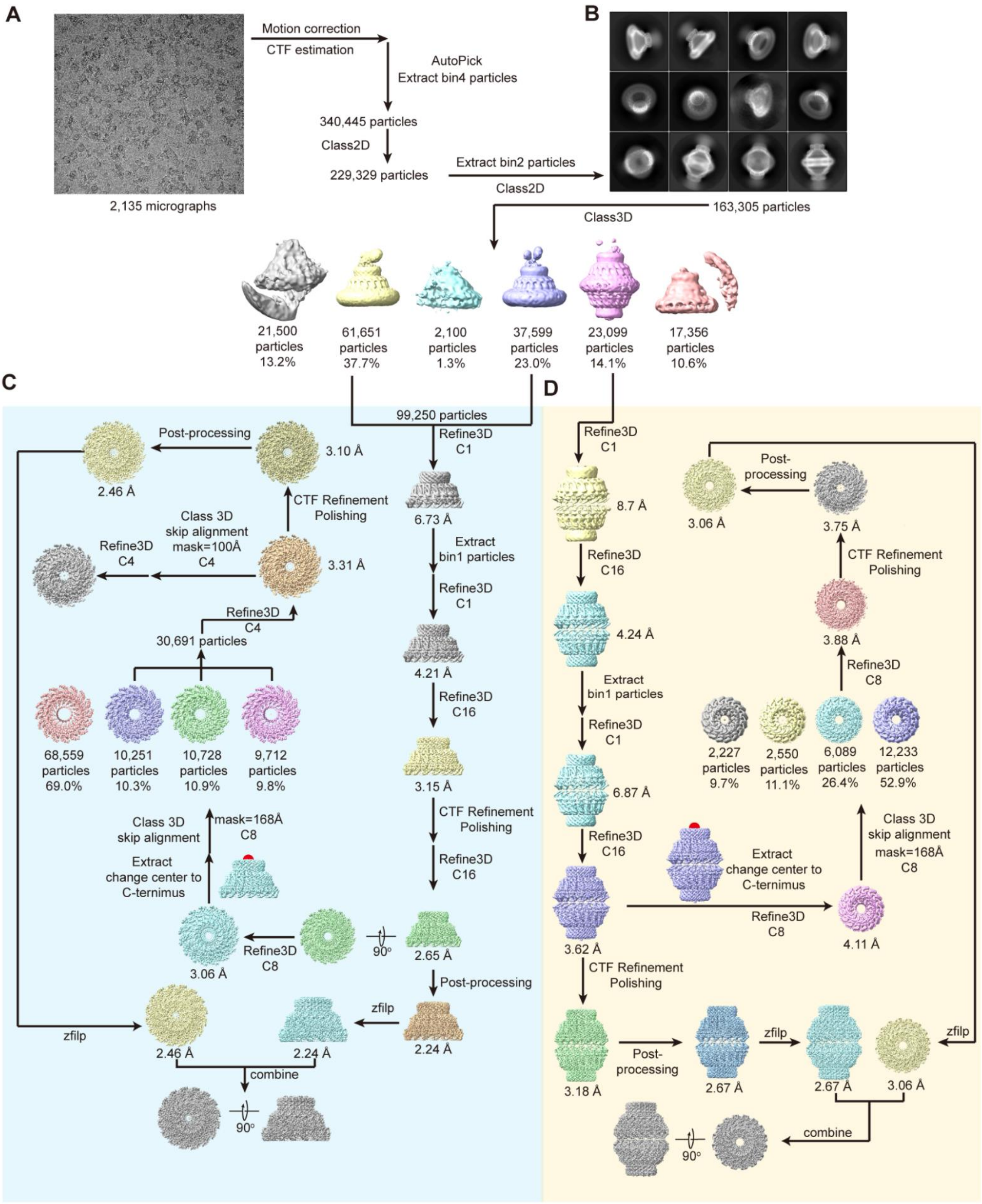
A
Motion correction
CTF estimation
AutoPick
Extract bin4 particles
340,445 particles
Class2D
229,329 particles
Extract bin2 particles
Class2D
2,135 micrographs
B
163,305 particles
Class3D
21,500 particles 13.2%
61,651 particles 37.7%
2,100 particles 1.3%
37,599 particles 23.0%
23,099 particles 14.1%
17,356 particles 10.6%
C
99,250 particles
Refine3D C1
6.73 Å
Extract bin1 particles
Refine3D C1
4.21 Å
Refine3D C16
3.15 Å
CTF Refinement Polishing
Refine3D C16
2.65 Å
90°
Post-processing
2.24 Å
zfilp
2.24 Å
Refine3D C8
3.06 Å
Extract change center to C-ternimus
Class 3D skip alignment
mask=168Å C8
68,559 particles 69.0%
10,251 particles 10.3%
10,728 particles 10.9%
9,712 particles 9.8%
30,691 particles
Refine3D C4
3.31 Å
Class 3D skip alignment mask=100Å C4
Refine3D C4
CTF Refinement Polishing
3.10 Å
Post-processing
2.46 Å
zfilp
2.46 Å
combine
90°
D
Refine3D C1
8.7 Å
Refine3D C16
4.24 Å
Extract bin1 particles
Refine3D C1
6.87 Å
Refine3D C16
3.62 Å
Extract change center to C-ternimus
Refine3D C8
4.11 Å
Class 3D skip alignment mask=168Å C8
2,227 particles 9.7%
2,550 particles 11.1%
6,089 particles 26.4%
12,233 particles 52.9%
Refine3D C8
3.88 Å
CTF Refinement Polishing
3.75 Å
Post-processing
3.06 Å
CTF Refinement Polishing
3.18 Å
Post-processing
2.67 Å
zfilp
2.67 Å
3.06 Å
zfilp
combine
90°

**Figure S2. Data processing of stomatin purified by DDM.**

**(A)** Representative raw cryo-EM micrographs of stomatin particles. A total of 2,135 micrographs were collected.

**(B)** Automated particle picking using the Laplacian-of-Gaussian algorithm identified 340,445 particles. After multiple rounds of 2D and 3D classification under bin4 and bin2, classes displaying well-defined single-layer and double-layer structures were selected. Ultimately, 99,250 particles (~60%) were retained for single-layer reconstruction and 23,099 (~14%) for double-layer reconstruction.

**(C)** Workflow for single-layer reconstruction. The 99,250 single-layer particles were subjected to 3D classification (bin2). Initial refinement yielded a 6.73 Å map, which revealed a clear 16-mer assembly. Reconstruction with C16 symmetry improved the resolution to 3.15 Å. Subsequent CTF refinement, Bayesian polishing, and post-processing yielded a final high-resolution map at 2.24 Å. The C-terminal density was further improved by mask-based 3D classification (skip alignment, C8 symmetry), resulting in a 2.46 Å map.

**(D)** Workflow for double-layer reconstruction. The 23,099 selected particles were used for 3D reconstruction. After CTF refinement, Bayesian polishing, and post-processing, a final map at 2.67 Å resolution was obtained. The C-terminal density was further improved by mask-based 3D classification (skip alignment, C8 symmetry), resulting in a 3.06 Å map.

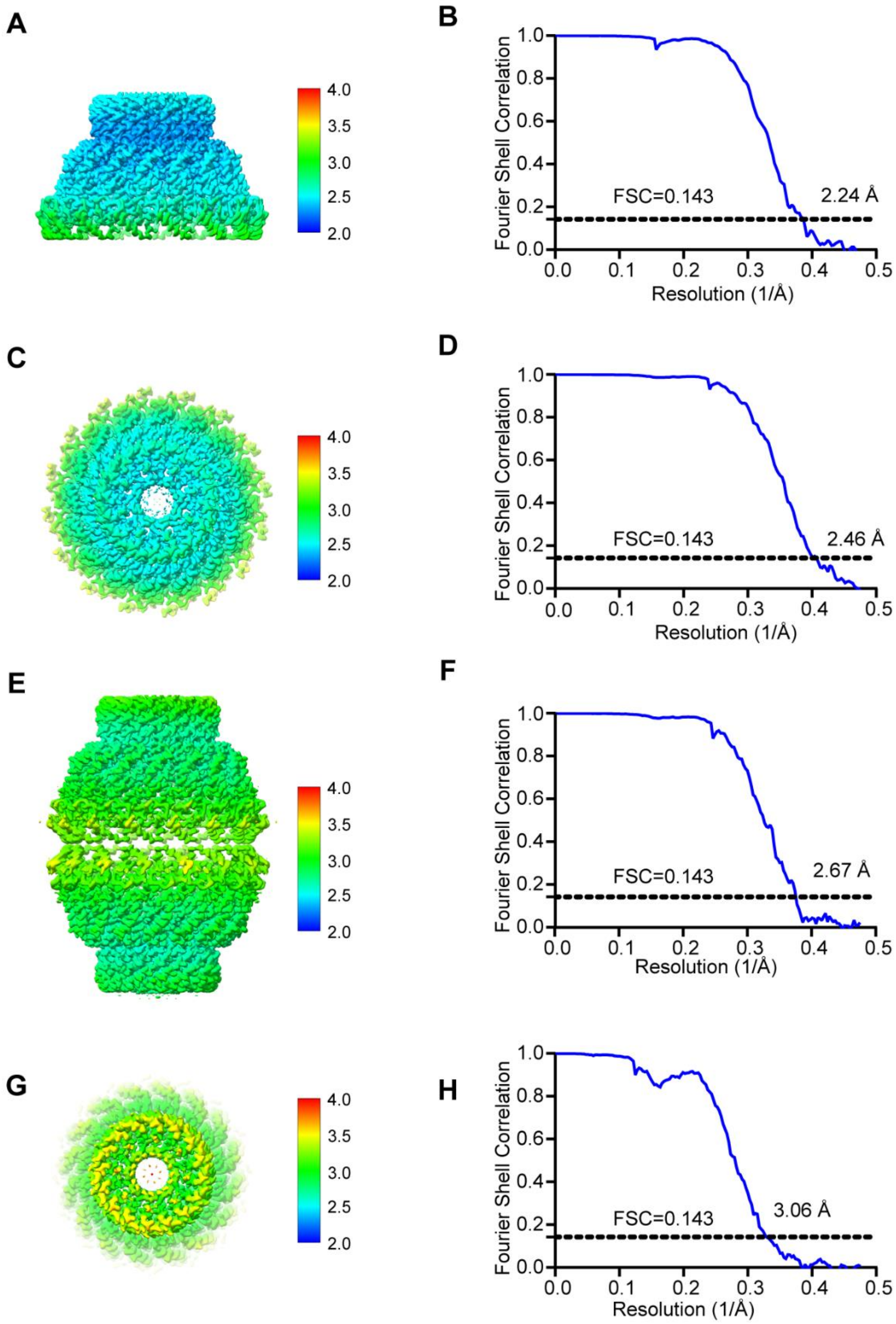

**Figure S3. Resolution estimation of reconstructed density maps of stomatin.**

Local resolution maps and Fourier shell correlation (FSC) curves of stomatin reconstructed under DDM solubilization conditions.

**(A–B)** Overall structure of STOM-single.

**(C–D)** C-terminal region of STOM-single.

**(E–F)** Overall structure of STOM-double.

**(G–H)** C-terminal region of STOM-double.

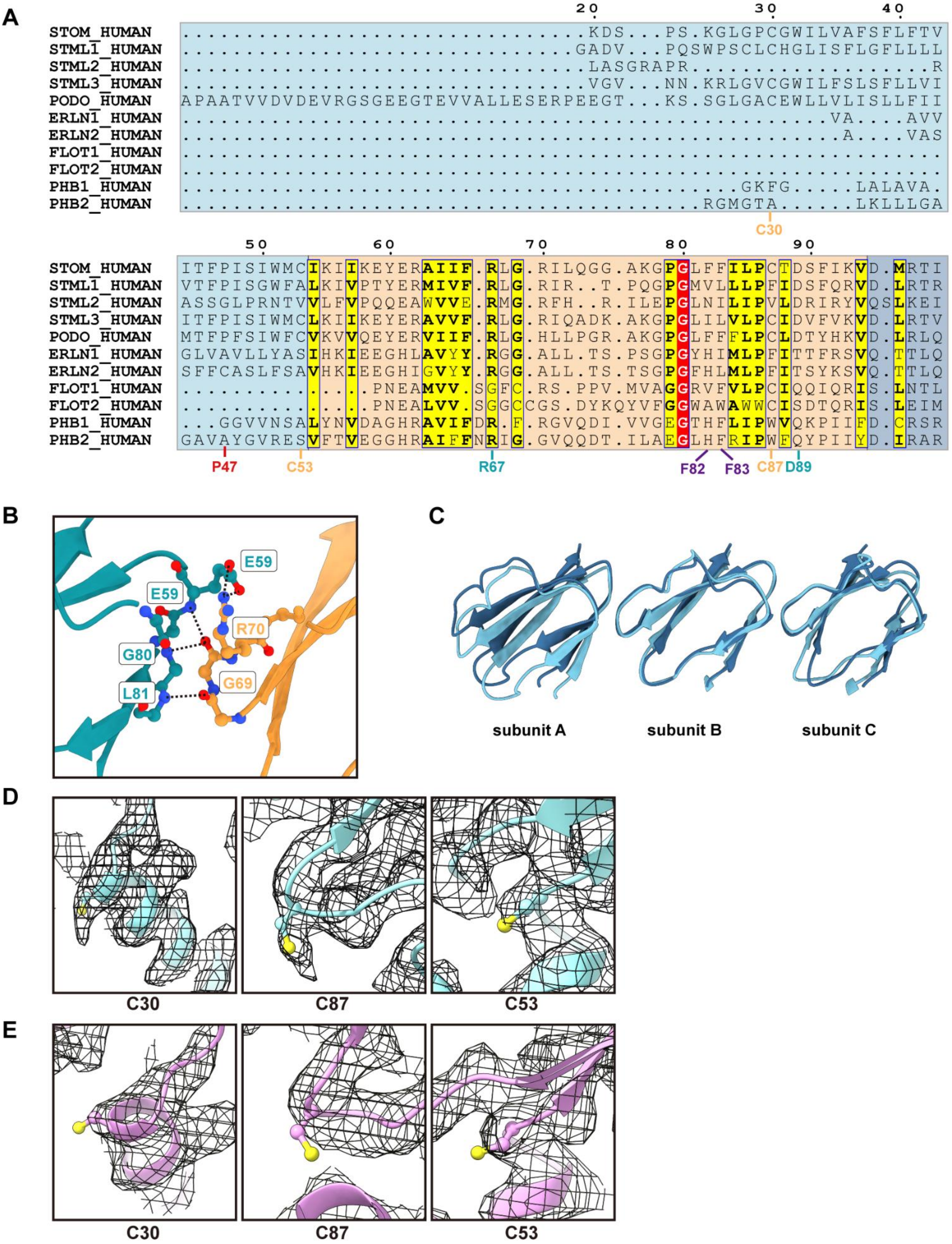
A
20
30
40
STOM_HUMAN .............................KDS...PS.KGLGPCGWILVAFSFLFTV
STML1_HUMAN ............................GADV...PQSWPSCLCHGLISFLGFLLLL
STML2_HUMAN .............................LASGRAPR...................R
STML3_HUMAN .............................VGV...NN.KRLGVCGWILFSLSFLLVI
PODO_HUMAN APAATVVDVDEVRGSGEEGTEVVALLESERPEEGT...KS.SGLGACEWLLVLISLLFII
ERLN1_HUMAN ..................................................VA....AVV
ERLN2_HUMAN ...................................................A....VAS
FLOT1_HUMAN .........................................................
FLOT2_HUMAN .........................................................
PHB1_HUMAN ...........................................GKFG.....LALAVA.
PHB2_HUMAN ........................................RGMGTA.....LKLLLGA
C30
50
60
70
80
90
STOM_HUMAN ITFPISIWMCIKIIKEYERAIIF.RLG.RILQGG.AKGPGLFFILPCTDSFIKVD.MRTI
STML1_HUMAN VTFPISGWFALKIVPTYERMIVF.RLG.RIR..T.PQGPGMVLLLPFIDSFQRVD.LRTR
STML2_HUMAN ASSGLPRNTVVLFVPQQEAWVVE.RMG.RFH..R.ILEPGLNILIPVLDRIRYVQSLKEI
STML3_HUMAN ITFPISIWMCLKIIKEYERAVVF.RLG.RIQADK.AKGPGLILVLPCIDVFVKVD.LRTV
PODO_HUMAN MTFPFSIWFCVKVVQEYERVIIF.RLG.HLLPGR.AKGPGLFFFLPCLDTYHKVD.LRLQ
ERLN1_HUMAN GLVAVLLYASIHKIEEGHLAVYY.RGG.ALL.TS.PSGPGYHIMLPFITTFRSVQ.TTLQ
ERLN2_HUMAN SFFCASLFSAVHKIEEGHIGVYY.RGG.ALL.TS.TSGPGFHLMLPFITSYKSVQ.TTLQ
FLOT1_HUMAN ...............PNEAMVV.SGFC.RS.PPV.MVAGGRVFVLPCIQQIQRIS.LNTL
FLOT2_HUMAN ...............PNEALVV.SGGCCGS.DYKQYVFGGWAWAWWCISDTQRIS.LEIM
PHB1_HUMAN ...GGVVNSALYNVDAGHRAVIFDR.F.RGVQDI.VVGEGTHFLIPWVQKPIIFD.CRSR
PHB2_HUMAN GAVAYGVRESVFTVVEGGHRAIFFNRIG.GVQQDT.ILAEGLHFRIPWFQYPIIYD.IRAR
P47
C53
R67
F82
F83
C87
D89
B
E59
E59
R70
G80
G69
L81
C
subunit A
subunit B
subunit C
D
C30
C87
C53
E
C30
C87
C53

**Figure S4. Structural features of the membrane-inserting region and lipid interactions of stomatin.**

**(A)** Sequence alignment of the NTD and SPFH1 domains of SPFH family proteins. Background colors indicate domain organization: NTD, light blue; SPFH1, orange; SPFH2, dark blue.

**(B)** Polar SPFH1–SPFH1 interface, formed primarily through loops connecting β-barrels.

**(C)** Alignment and comparison of SPFH1 domain in STOM-single (dark blue) and STOM-double (light blue).

**(D–E)** Cryo-EM maps showing densities of cysteine residues C30, C53, and C87 in STOM-single (D) and STOM-double (E).

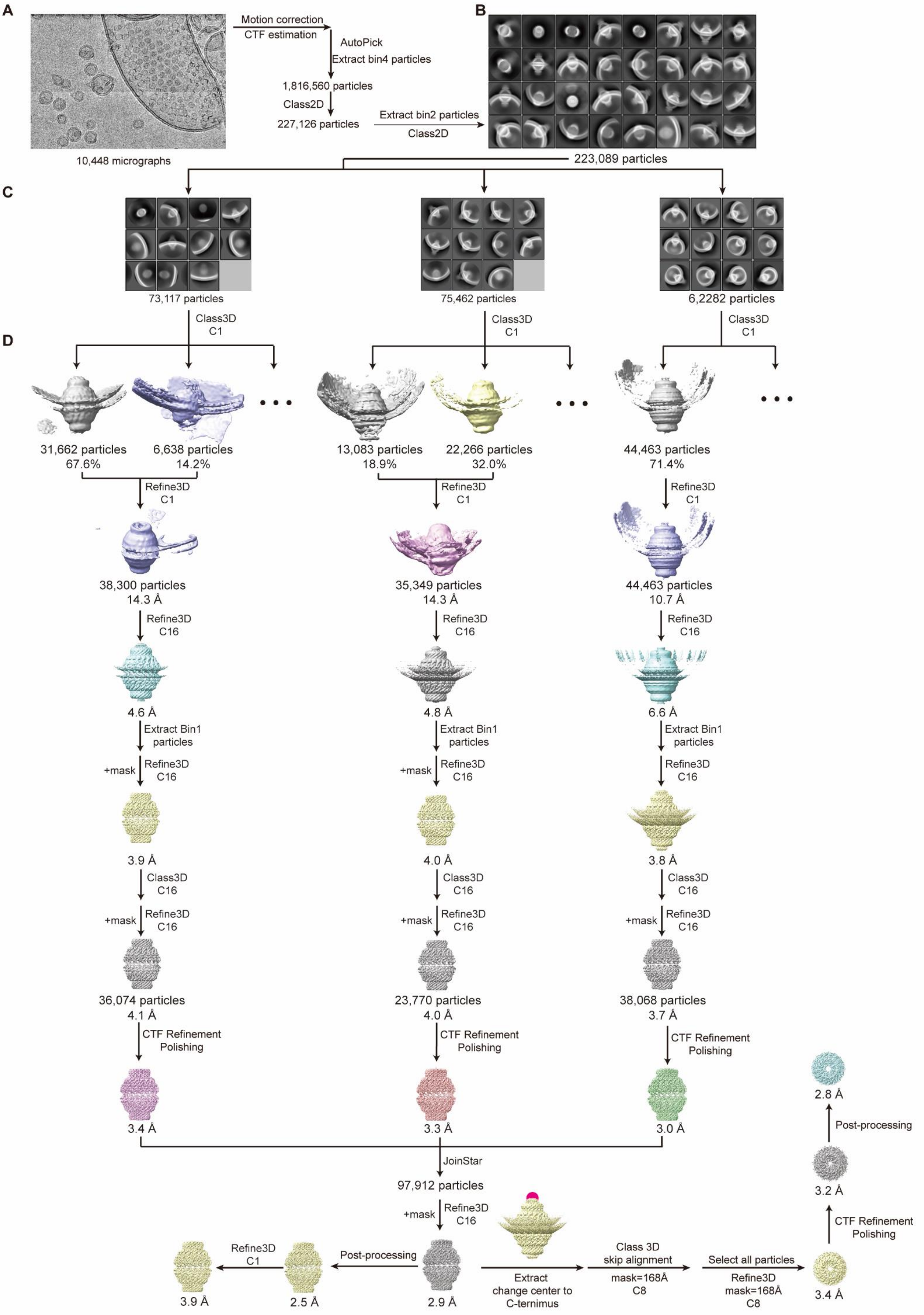
A
Motion correction
CTF estimation
AutoPick
Extract bin4 particles
1,816,560 particles
Class2D
227,126 particles
Extract bin2 particles
Class2D
10,448 micrographs
B
223,089 particles
C
73,117 particles
75,462 particles
6,2282 particles
Class3D
C1
D
31,662 particles
67.6%
6,638 particles
14.2%
13,083 particles
18.9%
22,266 particles
32.0%
44,463 particles
71.4%
Refine3D
C1
38,300 particles
14.3 Å
35,349 particles
14.3 Å
44,463 particles
10.7 Å
Refine3D
C16
4.6 Å
4.8 Å
6.6 Å
Extract Bin1
particles
+mask
Refine3D
C16
3.9 Å
4.0 Å
3.8 Å
Class3D
C16
36,074 particles
4.1 Å
23,770 particles
4.0 Å
38,068 particles
3.7 Å
CTF Refinement
Polishing
3.4 Å
3.3 Å
3.0 Å
JoinStar
97,912 particles
2.9 Å
Post-processing
2.5 Å
3.9 Å
Extract
change center to
C-ternimus
Class 3D
skip alignment
mask=168Å
C8
Select all particles
Refine3D
mask=168Å
C8
3.4 Å
3.2 Å
2.8 Å

**Figure S5. Data processing of STOM reconstruction on liposomes.**

**(A)** Representative raw cryo-EM micrographs of stomatin particles. A total of 10,448 micrographs were collected.

**(B)** Automated particle picking using the Laplacian-of-Gaussian algorithm identified 1,816,560 particles. After multiple rounds of 2D classification (bin4 and bin2), 223,089 particles were retained for 3D classification.

**(C)** Vesicles were grouped according to diameter.

**(D)** Stomatin particles from vesicles of different diameters were reconstructed separately, yielding maps at 3.4 Å, 3.3 Å, and 3.0 Å resolution. Merging these datasets produced a 2.9 Å structure. Subsequent CTF refinement, Bayesian polishing, and post-processing improved the map to 2.5 Å resolution. Mask-based 3D classification of the C-terminal region (skip alignment, C8 symmetry) further enhanced local density, yielding a 2.8 Å map.

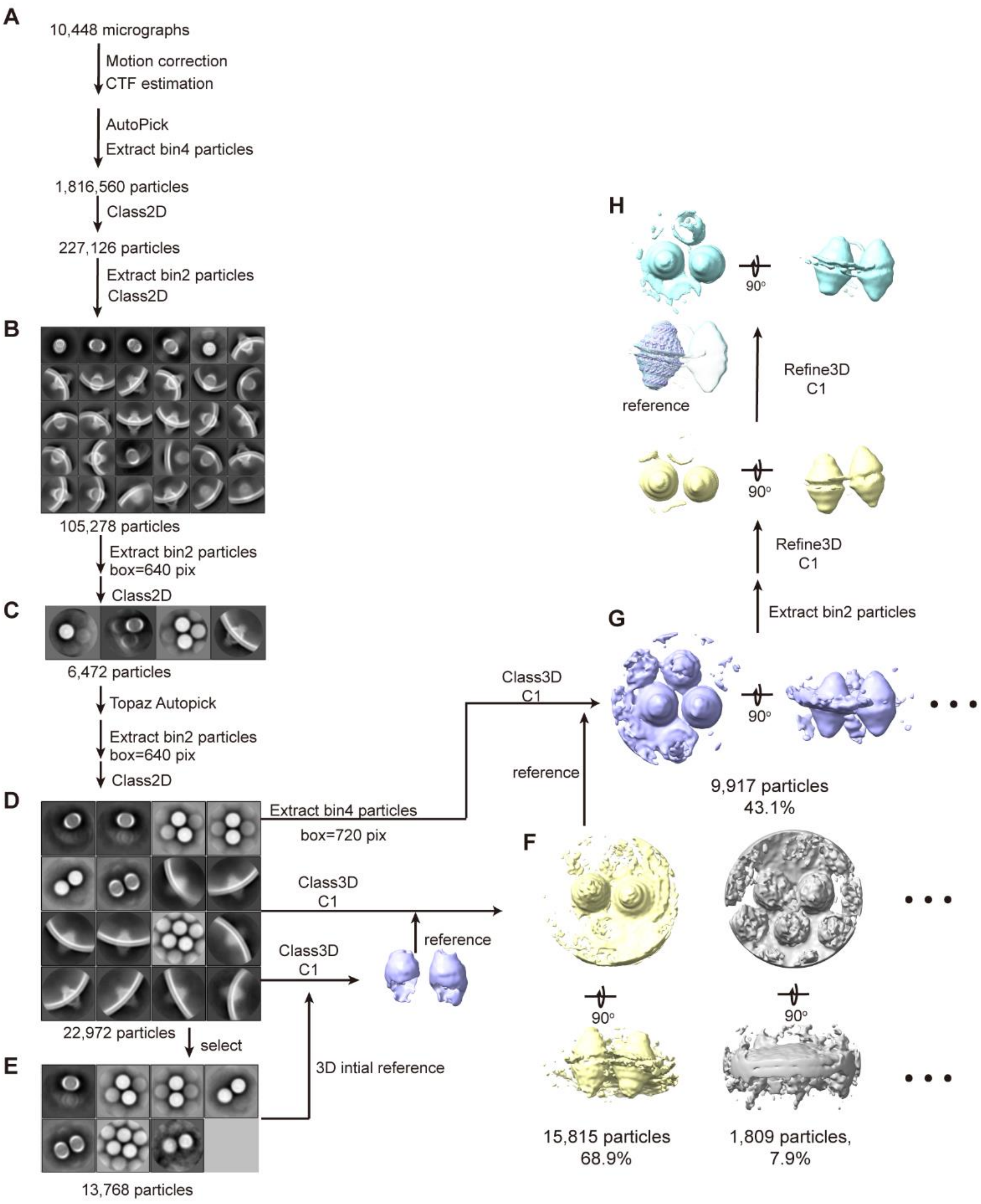
A
10,448 micrographs
Motion correction
CTF estimation
AutoPick
Extract bin4 particles
1,816,560 particles
Class2D
227,126 particles
Extract bin2 particles
Class2D
B
105,278 particles
Extract bin2 particles
box=640 pix
Class2D
C
6,472 particles
Topaz Autopick
Extract bin2 particles
box=640 pix
Class2D
D
22,972 particles
select
E
13,768 particles
Extract bin4 particles
box=720 pix
Class3D
C1
Class3D
C1
reference
3D intial reference
Class3D
C1
reference
F
15,815 particles
68.9%
1,809 particles,
7.9%
90°
G
9,917 particles
43.1%
90°
Extract bin2 particles
Refine3D
C1
90°
Refine3D
C1
reference
H
90°

**Figure S6. Data processing of STOM-cluster reconstruction on liposomes.**

**(A)** Initial processing followed the same workflow as in Figure S5.

**(B)** After multiple rounds of 2D classification, vesicle particles with the smallest curvature were selected.

**(C–D)** Re-extraction with a 640-pixel box followed by 2D classification revealed clear clustering of stomatin.

**(E)** The class displaying the most pronounced clustering was used to generate the initial model for 3D classification.

**(F)** The class with the clearest adjacent particle density was selected for iterative refinement.

**(G)** Particles in bin4 yielded the best classification quality.

**(H)** A total of 9,917 particles were used for 3D refinement, resulting in a low-resolution structure of stomatin cluster distribution.

### Supplementary Table 1

Cryo-EM data collection, refinement and validation statistics of stomatin

| | STOM-Single (EMD-XXXX) (PDB: YYYY) | STOM-double (EMD-XXXX) (PDB: YYYY) | STOM-liposome (EMD-XXXX) (PDB: YYYY) |
|---|---|---|---|
| **Data collection and processing** | | | |
| Electron microscope | Titan Krios | Titan Krios | Krios G4 |
| Electron detector | Gatan K2 | Gatan K2 | Falcon 4 |
| Magnification | 130,000× | 130,000× | 105,000× |
| Voltage (kV) | 300 | 300 | 300 |
| Electron exposure ($e^{-}/Å^{2}$) | 64 | 64 | 60 |
| Defocus range (μm) | -0.7 to -1.3 | -0.7 to -1.3 | -1.2 to -1.8 |
| Pixel size (Å) | 1.052 | 1.052 | 0.95 |
| Micrographs | 2,135 | 2,135 | 10,448 |
| Symmetry imposed | C8 | C8 | C8 |
| Final particle images (total) | 99K | 23K | 90K |
| Map resolution (total) (Å) | 2.2 | 2.5 | 2.5 |
| FSC threshold | 0.143 | 0.143 | 0.143 |
| **Refinement** | | | |
| Model composition | | | |
| Non-hydrogen atoms | 32080 | 66944 | 66048 |
| Protein residues | 4096 | 8544 | 8544 |
| Ligands | CLR:16 | CLR: 32 | 0 |
| R.m.s. deviations | | | |
| Bond lengths (Å) | 0.003 | 0.003 | 0.003 |
| Bond angles (°) | 0.628 | 0.552 | 0.570 |
| Validation | | | |
| MolProbity score | 1.65 | 1.65 | 1.20 |
| Clashscore | 6.59 | 6.78 | 2.92 |
| Rotamer outliers (%) | 2.35 | 1.73 | 1.41 |
| Ramachandran plot | | | |
| Favored (%) | 98.11 | 97.51 | 98.12 |
| Allowed (%) | 1.89 | 2.49 | 1.88 |
| Outliers (%) | 0.00 | 0.00 | 0.00 |
